# Thermally activated irreversible homogenization of G-quadruplexes in an ALS/FTD-associated gene

**DOI:** 10.1101/2025.06.02.657482

**Authors:** Daniel Ross, Olivia Lewis, Olivia McLean, Sundeep Bhanot, Shane Donahue, Rachael Baker, Randi Dias, David Eagerton, Vaibhav Mohanty, Bidyut K. Mohanty

**Affiliations:** Edward Via College of Osteopathic Medicine–Carolinas Campus, Spartanburg, SC 29303; Department of Chemistry and Chemical Biology, Harvard University, Cambridge, MA 02138; Harvard/MIT MD-PhD Program, Harvard Medical School, Boston, MA 02115 and Massachusetts Institute of Technology, Cambridge, MA 02139; Program in Health Sciences and Technology, Harvard Medical School, Boston, MA 02115 and Massachusetts Institute of Technology, Cambridge, MA 02139

## Abstract

A significant proportion of familial amyotrophic lateral sclerosis (ALS) and frontotemporal dementia (FTD) cases exhibit a substantial copy number expansion of the hexanucleotide GGGGCC/GGCCCC sequence in the *C9ORF72* gene. The GGGGCC sequence forms a non-canonical DNA structure called a G-quadruplex (G4) which has been associated with the disease states and with nucleic acid condensate formation. G4s can fold into various topologies, which can differentially impact fidelity of DNA synthesis. However, how G4 conformational heterogeneity and its regulation impact hexanucleotide repeat expansion is unclear, and important clues may lie in the thermodynamic properties of different G4 topologies. Here, we use temperature-swept CD spectroscopy to observe configurational homogenization of an initially heterogeneous population of G4s over a small range of temperatures, demonstrating thermally activated behavior. We further show that this reaction is irreversible, since subsequent temperature sweeps do not show CD shifts from non-parallel to parallel G4 topologies. Finally, we provide an analytical theory based on a two-state thermodynamic model which is compatible with experimental evidence, and we discuss alternate mechanisms for the homogenization transition. These findings suggest that kinetic regulation of non-canonical DNA structures may play a role in cellular homeostasis or disease pathogenesis.

**SIGNIFICANCE:** The GGGGCC repeats in the *C9ORF72* gene expand in copy number in certain neurodegenerative diseases, forming non-canonical DNA structures called G-quadruplexes (G4) which are associated with the pathological state. However, why the repeat expansion occurs is not known, and a key may lie in the thermodynamic stability of certain G4 conformations. Here, we use CD spectroscopy to experimentally report thermally activated heat-induced G4 conformation homogenization, from a heterogeneous population to the parallel configuration. We derive an analytical biophysical theory which is compatible with this experimental observation, which is shown to be irreversible. Our *in vitro* tuning of the free energy landscape that modulates G4 conformational fidelity motivates a search for possible *in vivo* enzymatic regulators.

## 1 INTRODUCTION

Amyotrophic lateral sclerosis (ALS, Lou Gehrig’s disease) and frontotemporal dementia (FTD) are devastating and generally fatal neuromotor diseases that, respectively, cause muscle weakness and brain atrophy, substantially increasing morbidity and mortality of affected individuals (1–5). Some of the shared symptoms of ALS and FTD are changes in behavior, personality, and language, and persons with a spectrum of behavioral disturbances common to ALS and FTD form a clinical subtype known as ALS-FTD (5, 6). Presently, there are few treatments for these disorders, which at best may delay some of the disease symptoms (3, 6, 7).

ALS, FTD, and ALS-FTD can be sporadic or familial. Various genetic mutations have been identified in patients with familial ALS, FTD, or ALS-FTD, many of which involve mutations in the *C9ORF72* gene (2, 6–9). Intron 1 of *C9ORF72* typically contains 2-22 copies of the hexanucleotide repeat GGGGCC/GGCCCC which can expand to 700-1600 copies in pathological settings (5, 8, 9). Transcription and down-stream translation of these hexanucleotide repeat expansions (HREs) are key contributors to the pathogenesis of *C9ORF72*-associated ALS/FTD (10–12). Although current research on development of ALS/FTD therapeutics often focuses on RNA and translated protein toxicity, the origin of the HRE and ALS/FTD lies solely in DNA and therefore targeting the mechanism of HRE is of equal importance (3).

Although the *C9ORF72* gene was discovered in 2011 (9), the core mechanism of its HRE still remains unexplained. An important clue to HRE may reside in the structural organization of hexanucleotide repeats as the GGGGCC sequence forms the non-canonical DNA structure called a G-quadruplex (G4) (13–17), and its complimentary sequence, GGCCCC, forms an intercalating motif (i-motif) (18, 19). G4s can fold into different topologies, and it has been shown recently that these topologies can differentially impact fidelity of DNA synthesis (20). Investigating the relationship between HREs and G4 topologies first motivates experimental mechanisms for modulating G4 configurational composition. We have recently shown that various environmental factors including pH and temperature regulate G4 formation in this ALS/FTD-associated HRE, as well as other G4-forming sequences including those found in *BCL2, c-MYC, EGFR, HIF1-α, hRAS*, and *VEGF* (21). Of these sequences, we found that only the GGGGCC hexanucleotide repeat’s CD spectrum undergoes major changes in the locations of its dominant peak(s) due to a heating cycle, suggesting that temperature regulates G4 configurational composition. These findings have motivated deeper investigation of the temperature-dependent changes to the G4 spectra in the ALS/FTD-associated HRE, which we perform here.

Recent work on other G4-forming sequences has indicated that the antiparallel configuration can be an intermediate in the process of folding to the more stable parallel G4 form, either existing transiently (22–24) or acting as a rapidly formed metastable state which slowly converts to a parallel G4 (25) over the course of minutes. We find the ALS/FTD-associated GGGGCC HRE, on the other hand, to be stable at room temperature indefinitely unless heated. Current computational and experimental evidence suggests that antiparallel G4s are generally separated from parallel G4s by a transition state barrier which leads to slow kinetics (25). We experimentally manipulate the G4 free energy landscape through heating; signatures of metastability and kinetic trapping phenomena would include configurational changes in G4 due to heating which are irreversible.

In the present work, we combine theoretical and experimental approaches to probe the thermally activated transition of G4 configurations. We perform temperature-swept CD spectroscopy to obtain “3D CD” spectra (26, 27) showing how CD changes as a function of both wavelength and temperature. From these spectra, we observe the transition from a configurationally heterogeneous population of G4s into a homogeneous population of parallel G4s, followed by melting of the parallel G4s at even higher temperatures. We then perform sequential temperature sweeps with a cooling period in between to demonstrate the irreversibility of the transition. We finally introduce a simple analytical theory using a two-state model for G4 configuration prevalence motivated by the computational findings of ref. (25) which is compatible with the spectra obtained from our experiments. We then discuss alternative mechanisms which could result in similar experimental findings. Our work opens the door to understanding how the relative fractions of different DNA G4 configurations may be stabilized or regulated; since G4s are known to facilitate intracellular RNA condensate formation in pathological disease states, these results carry implications for coupling between G4 configuration switches and disease-associated liquid-liquid phase separation of DNA (12, 28–30).

## 2. MATERIALS AND METHODS

### DNA oligonucleotides

DNA oligonucleotides used in these studies were obtained commercially from Millipore (Burlington, MA) and Eurofins. All oligonucleotides were purified by high-performance liquid chromatography, shipped dry, and reconstituted with TE (10 mM Tris and 1 mM EDTA) or nuclease free water (Invitrogen) to 100 pmol/*μ*L final concentration.

### Buffers

The buffers used in these studies are 10 mM sodium cacodylate (NaCac) and 100 mM KCl at pH 7.4.

### CD measurements

All CD measurements were carried out with a Jasco J-1500 CD spectrophotometer with a single-position Peltier-thermo cell holder. Samples were measured in 1 mm path length Starna cuvettes. For the 3D CD experiments, temperature sweeps were set to start at 20 °C and end at 100 °C, with a temperature gradient of 1 °C/min and a wait time of 30 seconds between measurements, which occurred at intervals of 2 °C. An additional experiment was conducted with a wait time of 300 seconds between measurements. The temperature gradient was paused during measurement. The starting condition was set to maintain the initial temperature within ± 0.1 °C for 10 seconds. Extended range temperature sweeps were conducted in polyethylene glycol (PEG) from 20 °C to 100 °C. For the irreversibility experiments, the first sweep was conducted either from 20 °C to 80 °C or from 20 °C to 100 °C, while the second sweep was always conducted from 20 °C to 100 °C. If a third sweep was conducted, it was also conducted from 20 °C to 100 °C.

## 3 RESULTS

### 3.1 Experimental temperature-swept CD spectra reveal thermally activated G4 homogenization, followed by melting

Before introducing experimental results, we first discuss the standard spectroscopic signatures of various G4 conformations. In Figure 1, we schematically depict the different categories of G4 conformations. In parallel G4s (PG4s), all four DNA strands are oriented in the same direction (Figure 1A). In antiparallel G4s, the four DNA strands are in alternating orientation, and in hybrid G4s, three out of four strands are oriented in one direction while one is oriented in the opposite direction. Parallel, antiparallel, and hybrid G4s have distinct CD peaks. Parallel G4s have a negative peak at 240 nm and a larger positive peak around 260 nm, antiparallel G4s have a negative peak at 260 nm and a positive peak at 295 nm, and hybrid G4s show positive peaks at both 260 nm and 295 nm and a negative peak around 240 nm (31). It is experimentally difficult to distinguish the CD spectrum of hybrid G4s from a mixed population of antiparallel and parallel G4s, so throughout this work we informally group together hybrid G4s and antiparallel G4s together as “non-parallel” G4s (NPG4s) (Figure 1B).

**Figure 1.**
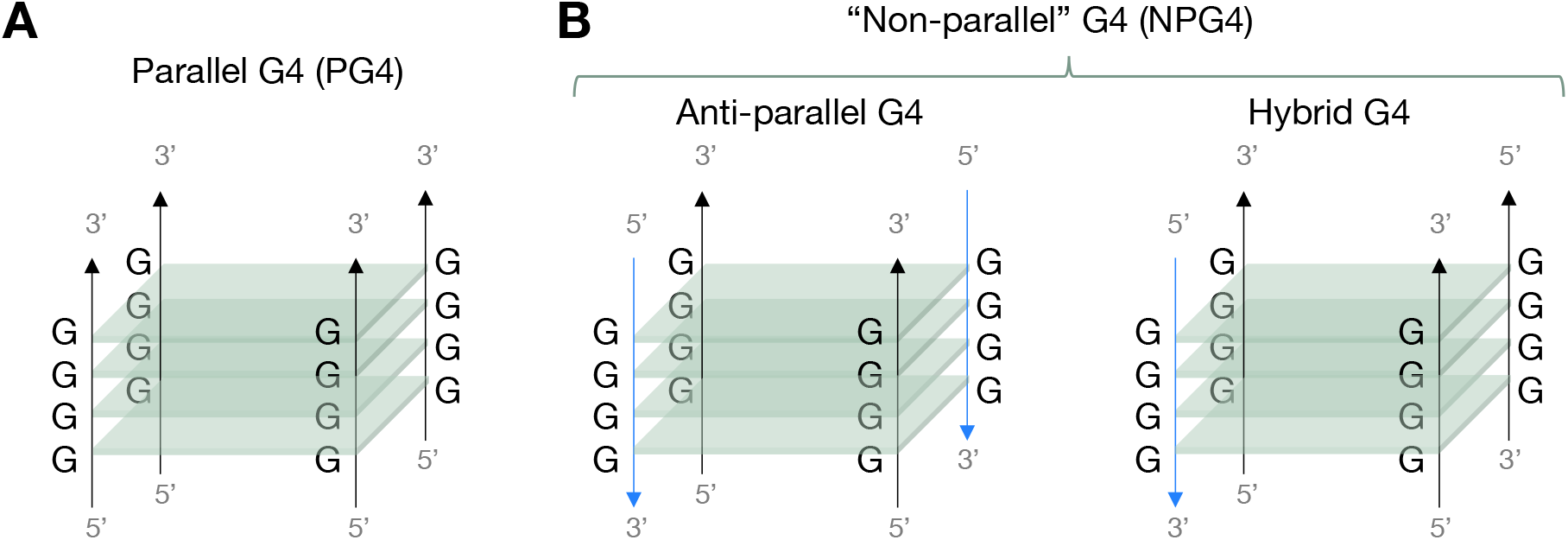
Schematic representation of G4 structures. (**A**) Parallel G4s (PG4s) have all 4 DNA strands parallel to each other. (**B**) Antiparallel G4s have their DNA strands alternating orientations, and hybrid G4s have three DNA strands parallel to one another and a fourth one antiparallel to the first three. We group antiparallel and hybrid into a category called “non-parallel” G4s (NPG4s).

To probe the configurational composition of G4 populations experimentally, we conducted temperature-swept CD spectroscopy of DNA oligonucleotides containing the GGGGCC hexanucleotide repeat. This involved taking CD spectra of the same DNA sample as the temperature was incrementally increased from 20 °C to 100 °C in increments of 2 °C, with 30 second wait periods before CD measurement. An extended range temperature sweep experiment was also conducted in the presence of PEG from 20 °C to 110 °C. The resulting data were 3D CD spectra which visualize circular dichroism as a surface plot against both wavelength and temperature. The same data is also visualized as a set of superimposed typical 2D spectra with a color gradient to indicate the temperature.

Experimental 3D CD spectra as well as superimposed standard 2D CD spectra with color-coded temperatures of hexanucleotide repeats with various copy numbers are plotted in Figure 2. For 4 repeats, 14 repeats, and 20 repeats (as well as for 16 repeats in PEG), we observe that the initial samples at 20 °C show peaks that match hybrid or a mixed population of antiparallel, hybrid, and parallel G4s—an indication that we likely have a configurationally heterogeneous population of G4s (Figure 2A-D) at room temperature, which agrees with findings in the literature which were based on NMR spectra (32). These initial spectra are stable for long times at room temperature; we found that for a 17-repeat oligomer, the CD spectrum remained essentially exactly the same every day for five days (Figure S1).

**Figure 2.**
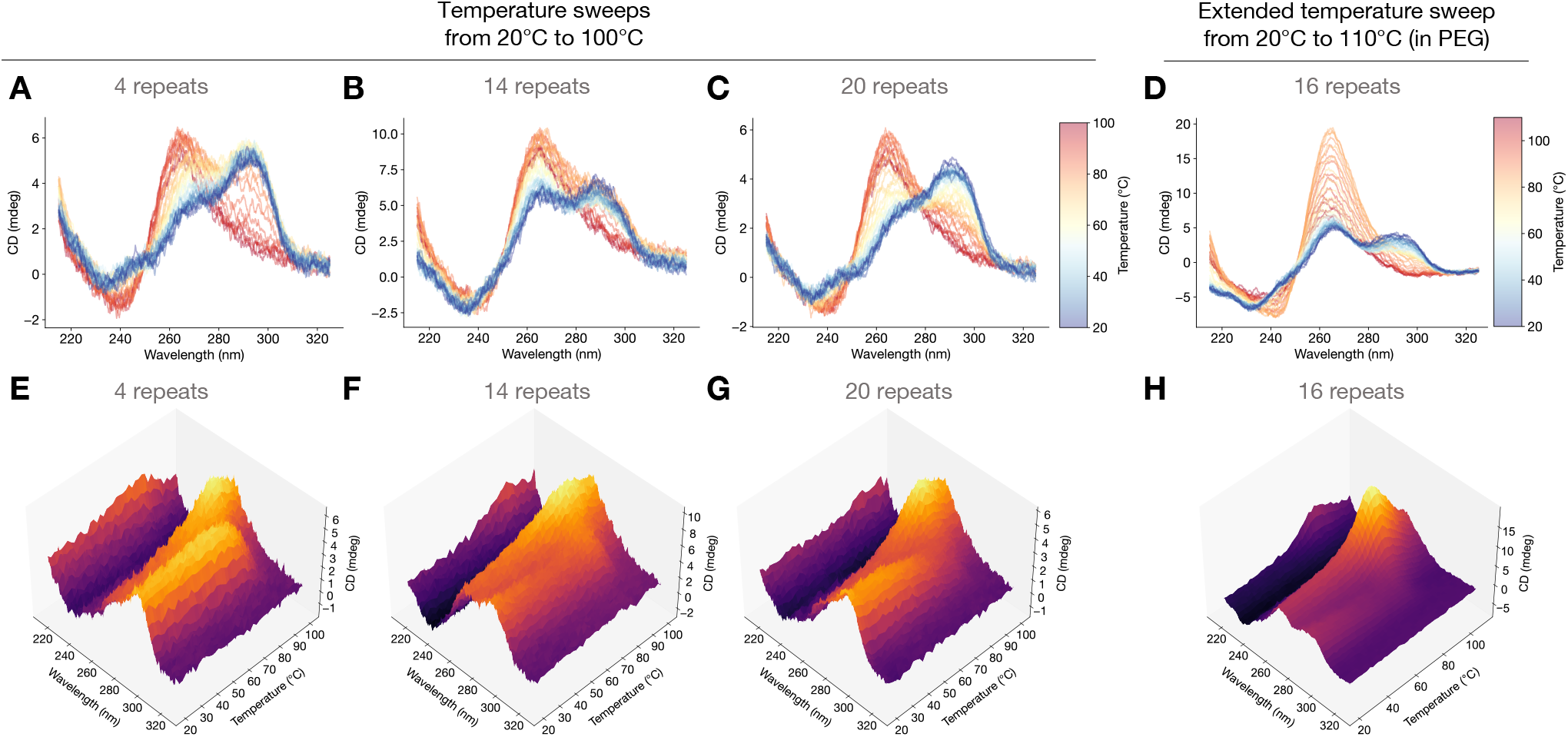
Temperature-swept CD spectra reveal thermally activated homogenization to PG4s followed by melting of PG4s at subsequently higher temperatures. (**A**,**B**,**C**) Temperature-swept CD spectra swept from 20 °C (blue) to 100 °C (red) for 4, 14, and 20 repeats, respectively. (**D**) Extended range temperature-swept CD spectrum swept from 20 °C (blue) to 110 °C (red) for 16 repeats, in PEG. (**E**,**F**,**G**,**H**) Same data as (A,B,C,D), but plotted as 3D CD spectra showing temperature-swept circular dichroism, revealing decreasing peaks around 290 nm and increasing peaks around 260 nm until some temperature *T*_peak_, above which the 260 nm peak decays.

The final CD spectra, at the maximum temperature of 100 °C (or 110 °C for the extended temperature experiment), all match the spectroscopic signature of parallel G4s, suggesting homogenization at some intermediate temperature (Figure 2A-D). Both the 2D and 3D CD spectra showing the continuous dependence on both wavelength and temperature indicate, that regardless of copy number, the 290 nm peak is stable from the starting temperature 20 °C up to roughly 70 °C, after which the 260 nm peak rapidly rises to prominence at higher temperatures (Figure 2A-H). The profile of the experimental 3D CD spectra is highly indicative that a configurational *homogenization* from NPG4s (or, more likely, a mixed starting population of NPG4s and PG4s) occurs over a small window of temperatures. A theoretical mechanism for this change is proposed in section section 3.3. We performed the same temperature-swept CD spectroscopy for additional copy number variants, from 2 repeats to 20 repeats, and received similar results (Figure S2, S3, S4, and S5; replicates and experiments with various repeat numbers summarized in Table S1). Moreover, to confirm that CD spectroscopic changes associated with the transition is not limited to oligonucleotides from one commercial source, we tested the effect of temperature increase on CD with oligonucleotides from another commercial source and obtained the same results.

Each of the temperature-swept CD spectra in Figure 2 display a characteristic increase of the ~ 260 nm peak until some temperature *T*_peak_ after which the ~ 260 nm peak begins to decay. This decay is especially apparent in the extended temperature sweep for 16 repeats (Figure 2D,H). The CD spectrum at *T*_peak_ possesses the characteristics of a typical parallel G4 spectrum, with a positive peak around 260 nm, a negative peak around 240 nm, and a smooth decay from the 260 nm peak toward zero for higher wavelengths (Figure 2A-D). For the extended temperature range sweep, we now examine the correlations between the spectrum at *T*_peak_ and the spectrum at other temperatures in order to better understand the conformational composition of the G4 population.

At temperatures *T* ranging from 20 °C to *T*_peak_ = 84 °C, there is initially a notable non-linearity in the plot between the CD spectrum at *T* versus the CD spectrum at *T*_peak_ which disappear as *T* approaches *T*_peak_ (Figure 3A). A straight line would indicate that the spectra take on the same shape but may have differing magnitude. Near room temperature, the CD spectrum at *T*, however, has deviations from a straight line which is indicative of differing configurational composition. Since NPG4s have different characteristic positive and negative peaks from PG4s, we can interpret the gradual progression of the non-linear spectrum-spectrum correlation into a linear one as reflecting a configurational change—particularly, one in which a heterogeneous G4 population ultimately becomes all PG4s, since the spectrum at *T*_peak_ = 84 °C possesses the characteristic peaks of a PG4 spectrum.

**Figure 3.**
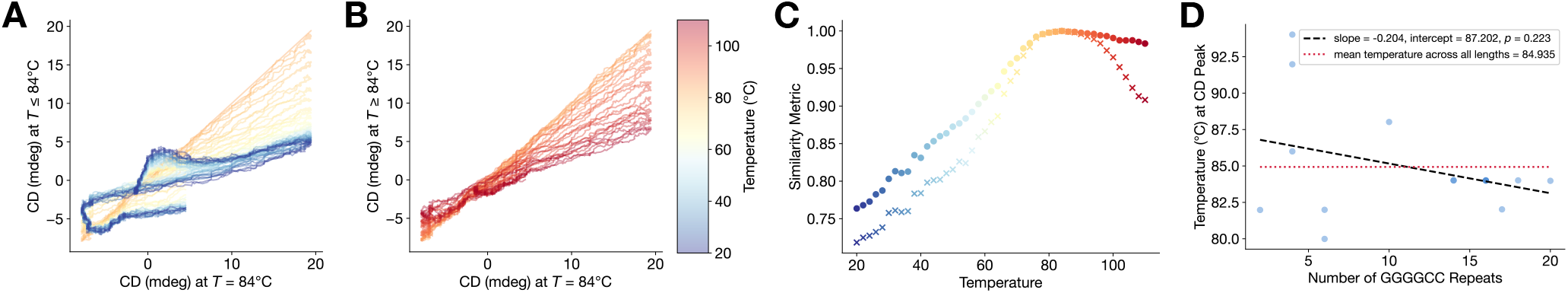
Spectrum-spectrum correlations reveal G4 compositions in different temperature regimes. (**A**) Plot of CD spectra at temperatures *T* ranging from 20 °C to *T*_peak_ = 84 °C versus CD spectrum at *T*_peak_ = 84 °C, showing configurational change. (**B**) Plot of CD spectra at temperatures *T* ranging from *T*_peak_ = 84 °C to 110 °C versus CD spectrum at *T*_peak_ = 84 °C, showing magnitude decay without spectral characteristic change, indicating melting. (**C**) Pearson correlation and cosine similarity between CD spectrum at temperature *T* and CD spectrum at temperature *T*_peak_. (**D**) *T*_peak_ versus repeat length across multiple replicates and experimental conditions, summarized in Table S1.

On the other hand, for *T* ≥ *T*_peak_ = 84 °C (until the maximum temperature of 110 °C) we observe that the spectrum-spectrum plots remain linear, but with decreasing slope (Figure 3B). This means that for temperatures *T* above *T*_peak_, the same CD spectrum *shape*—corresponding to PG4s—is maintained, but the *magnitude* decreases with increasing temperature. Since unfolded G4s have nearly flat spectra (33), this maintenance of spectral shape with decaying magnitude is indicative of PG4 melting as temperature is increased beyond *T*_peak_ = 84 °C. Estimates of melting temperature for GGGGCC oligomers in the literature are reported as “*T*_*m*_ > 84 °C, or even higher” (34) and “90 °C and 75 °C, respectively” for 4 and 5 hexanucleotide repeats (32), with ref. (14) noting incomplete “dissociation of the G-quadruplex even at 95 °C.” Interpreting Figure 3B as an indication that PG4s begin melting above 84 °C is thus consistent with reported melting temperatures in the literature.

The spectrum-spectrum correlation plots thus indicate two temperature regimes in which G4 composition changes: below *T*_peak_, a heterogeneous G4 population homogenizes into the parallel configuration. Then, above *T*_peak_, this homo-geneous population of PG4 melts as temperature is further increased. To quantify spectrum-spectrum correlations, we plot the Pearson correlation and cosine similarity between the CD spectrum at *T*_peak_ and the CD spectrum at the sweep temperature *T*. We see that Pearson correlation and cosine similarity increase nearly monotonically over the temperature range from 20 °C to *T*_peak_ = 84 °C. Above *T*_peak_, the cosine similarity decays to 0.908 at 110 °C—similar to its value of 0.916 at 66 °C—while the Pearson correlation hardly decays at all, retaining a value of 0.983 at 110 °C. This indicates that the CD spectra between *T*_peak_ and 110 °C are highly linearly correlated, consistent with out qualitative observations from Figure 3B, while this correlation is much weaker near room temperature, as observed in Figure 3A. Thus, our evidence supports the notion that G4 homogenization into PG4s occurs over the temperature sweep from room temperature to *T*_peak_ (which for 16-repeats in the presence of PEG is around 84 °C), and above this temperature the homogeneous PG4s unfold.

We then compute *T*_peak_ for many experimental trials with varying GGGGCC repeat lengths, ranging from 2 to 20, with multiple replicates for some oligomers. Our results, which include trials which include PEG or use extended wait times in the temperature gradient, are summarized in Table S1 and are plotted in Figure 3D. The plot shows no statistically significant linear relationship (*p*-value = 0.223) between *T*_peak_ and repeat length. Since we cannot reject the null hypothesis that *T*_peak_ and repeat length are uncorrelated, we calculate the mean of *T*_peak_ across repeat lengths as a summary statistic, finding that the average *T*_peak_ is 84.935 °C.

Lastly, we conducted the temperature sweep experiment for 4-repeat and 16-repeat oligomers, but with extended wait times at every point in the temperature gradient. Instead of incrementing the temperature by 2 °C with a wait time of 30 seconds between each increment, we waited 300 seconds between each increment. For 4-repeat and 16-repeat oligomers, the resulting CD spectra appear similar in both 2D and 3D representations (Figure S5), regardless of the wait time. The 4-repeat oligomers at room temperature in these wait time validation experiments have a different initial fraction of NPG4s and PG4s compared to the initial 4-repeat experiments presented in main text Figure 3A,E, possibly due to differences in company synthesis of oligomers or of sample preparation in the laboratory.

### 3.2 Irreversibility of G4 homogenization

We then explored if the apparent temperature-dependent homogenization to PG4s was reversible. A key signature of irreversibility would be if the homogeneous PG4 population were to remain homogeneous even if the system is cooled back to room temperature after the initial temperature sweep. A second or third temperature sweep would then lead to little change in the configurational composition, which means the CD spectrum would not change after the first sweep.

To investigate irreversibility experimentally, we sequentially performed a temperature sweep followed by a cooling period back to near room temperature followed by a second temperature sweep. For oligonucleotide samples with various copy numbers, we measured CD at a fixed wavelength corresponding to the principle positive peak for PG4s during each of the two temperature sweeps.

In Figure 4A, we show across three triplicates that for 6 hexanucleotide repeats that circular dichroism at 265 nm (corresponding to the PG4 parallel peak) generally increases with temperature during the sweep from 20 °C up to around *T*_peak_ ≈ 80 °C, consistent with previous section’s experimental results indicating a predominant presence of PG4s and a decline in NPG4s. Above *T*_peak_, the magnitude of the peak begins to decline until our maximum temperature of 100 °C. After reaching the maximum temperature, we allow the sample to cool back to room temperature. A second sweep then shows that the CD at room temperature *has not* returned to its original magnitude, but rather is closer in magnitude to the value reached at *T*_peak_. It generally stays flat until 60-70 °C, where the CD magnitude begins to increase again, peaking once again around the same *T*_peak_ ≈ 80 °C and subsequently declining. Across the three replicates, we noticed that the second sweep’s peak tended to be higher than the first sweep’s peak. The results in Figure 4A demonstrate clear sign of irreversibility of the homogenization transition between 20 °C and *T*_peak_ ≈ 80 °C, but the decline in the peak from *T*_peak_ ≈ 80 °C to 100 °C appears reversible, consistent with our analysis in the previous section.

**Figure 4.**
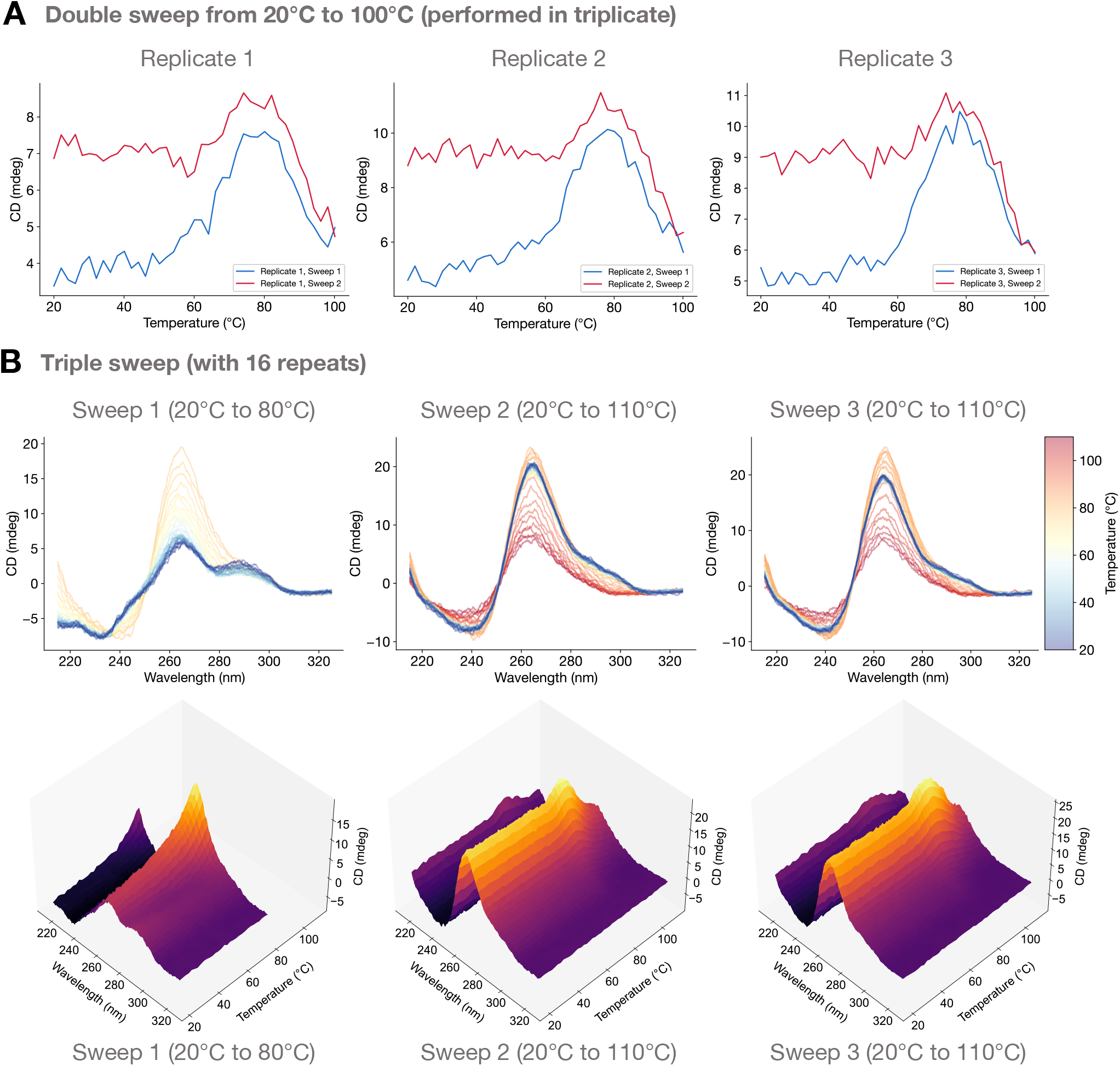
Thermally activated G4 homogenization is irreversible. (**A**) For 6 hexanucleotide repeats, the first temperature sweep (blue) shows a CD increase at around 265 nm as temperature increases from 20 °C up to 80 °C, indicative of a transition to PG4s, followed by a CD decrease above 80 °C due to melting. After cooling back to 20 °C, the second temperature sweep (red) shows stability of the CD peak from 20 °C up to roughly 80 °C, after which CD decreases, either due to G4 melting (or possibly NPG4 formation). Experiments conducted in triplicate. (**B**) For 16 hexanucleotide repeats, 3D CD spectra were measured over three temperature sweeps, first from 20 °C up to 80 °C, then subsequently from 20 °C up to 110 °C during the second and third sweeps. Irreversibility is clearly indicated by differences between the first and second sweeps, while the third sweep matches the second.

We then expanded beyond a single wavelength and conducted full 3D CD temperature sweeps (with accompanying 2D color-coded CD spectra), with an initial sweep from 20 °C to 80 °C followed by two additional sweeps from 20 °C to 110 °C (Figure 4B). We observed the characteristic disappearance of the antiparallel G4-associated positive peak at 290 nm during the first sweep with a simultaneous increase in the peak around 260 nm until reaching a peak value around 80 °C. The second and third sweeps then have qualitatively similar 3D CD spectra, with the entire spectrum remaining nearly constant across the entire temperature range from 20 °C up to around *T*_peak_ ≈ 80 °C, after which the spectrum decays in magnitude without changing morphological features, as observed in the previous section. Additional double-sweep irreversibility experiments conducted at fixed wavelength for 4, 14, and 20 repeats showcase similar results (Figure S6A-C). The irreversible nature of the thermally activated homogenization of G4 conformations suggests that metastability may play a role.

### 3.3 Compatible theoretical models of thermally activated G4 homogenization

In light of experimental data, we now develop a simple, minimal two-state analytical theory for the configurational state of G4s which is compatible with the presented experimental data. We subsequently discuss alternative mechanisms which may yield similar results. Mathematically, we will explicitly develop a theory which predicts a thermally activated metastable-to-stable transition, yielding a homogeneous population of parallel G4s due to heating. The theory yields simulated CD spectra which align with the experimentally observed CD spectra.

To summarize the two-state theory, we begin with a heuristic explanation. We expect that a heterogeneous population of NPG4s and PG4s are metastable for long times because of thermodynamic stability relative to the unfolded state, but they are separated by a tall transition state barrier which leads to very slow kinetics (Figure 5A). Since the transition state has higher entropy than the G4 states, increasing the temperature lowers the relative height of the free energy barrier to transition from NPG4s to PG4s, making the metastable-to-stable transition suddenly possible even at short timescales, sending the population to primarily the PG4 state (Figure 5B). Cooling the system then traps the population homogeneously in the thermodynamically stable PG4 state (Figure 5C). Further heating would keep the population predominantly in the PG4 state due to its thermodynamic stability relative to NPG4.

**Figure 5.**
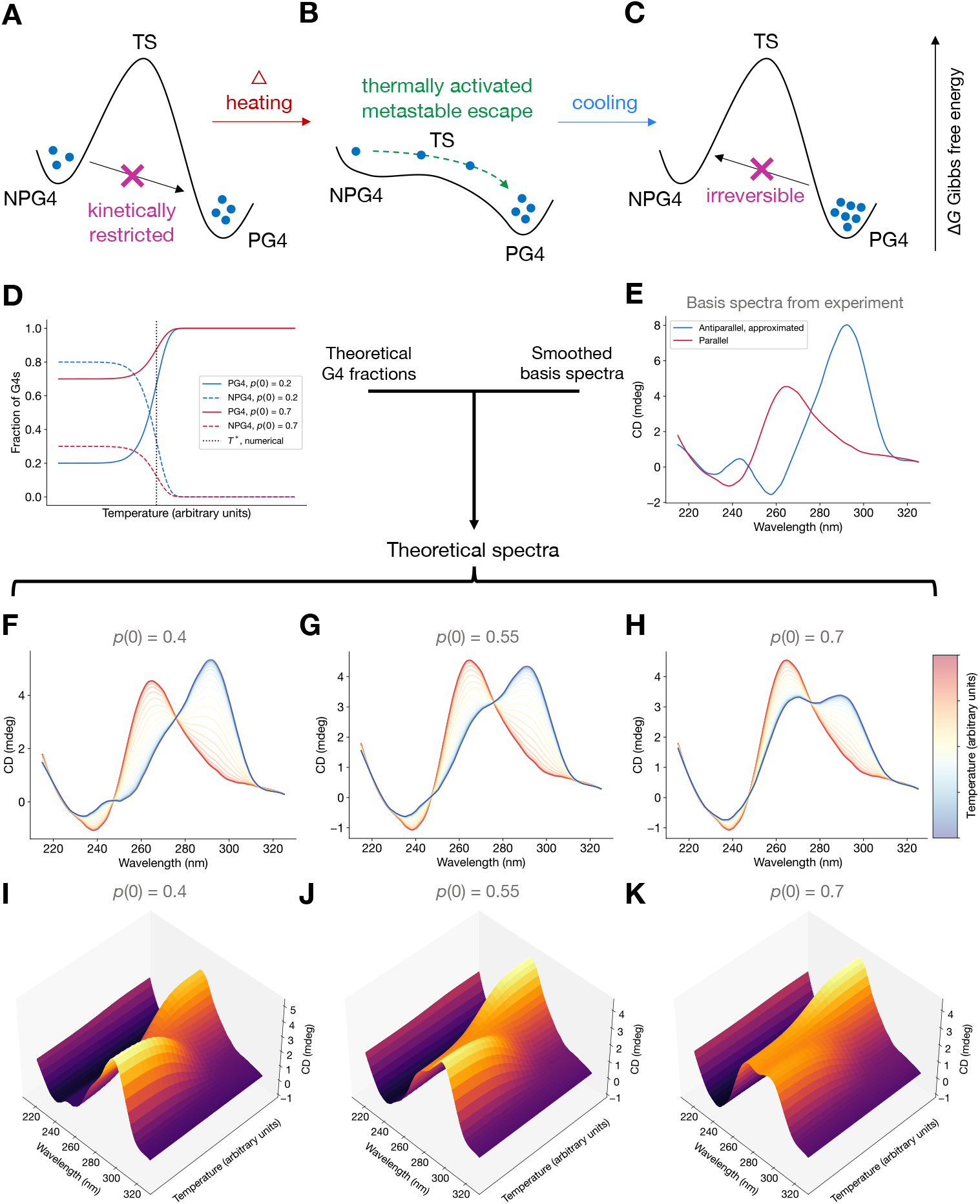
Compatible theoretical mechanism and simulated 2D and 3D CD spectra showcasing a thermally activated homogenization mechanism. (**A**) Initial heterogeneous population of G4s remain heterogeneous over long timescales due to transition state barrier separating NPG4s and PG4s. (**B**) A thermally activated metastable-to-stable transition: heating lowers the free energy barrier relative to the G4-forming states just enough to allow rapid transitions from the metastable NPG4 state to the stable PG4 state without unfolding G4s entirely. (**C**) Cooling the system once again imposes a transition state barrier for the reverse reaction for the now-homogeneous PG4 population. (**D**) Theoretical fraction of PG4s and NPG4s as a function of temperature indicates thermally activated behavior from a starting PG4 fraction *p* (0; *T*) and NPG4 fraction 1 − *p* (0; *T*) to a (nearly) homogeneous population of PG4s at higher temperatures. The homogenization temperature *T* ^*^, numerically computed as the inflection point, is plotted (black, dotted). Two starting fractions, *p* (0; *T*) = 0.2 (blue) and *p* (0; *T*) = 0.7 (red) are plotted, and *T* ^*^ was the same for both. (**E**) Experimentally determined PG4 basis CD spectrum *ϵ*_*P*_ (*λ*) (red) and approximated antiparallel basis CD spectrum *ϵ* _*A*_ (*λ*) (blue), used to construct theoretical 3D CD spectrum. (**F**,**G**,**H**) Theoretical temperature-dependent 2D CD spectra computed from eq. (9) by using eq. (7) and basis spectra from (E), plotted on an arbitrary temperature axis, reveals a CD positive peak shift from near 290 nm to near 265 nm, which is an indicator of NPG4s converting to PG4s. (**I**,**J**,**K**) Same as (F,G,H) but shown as 3D CD spectra.

To make the theory concrete, we first define some notation. We let *P* represent the PG4 state and let *N* represent the NPG4 state. These form the two discrete states in our thermodynamic model. We will use the double dagger ‡ superscript corresponds to the unfolded state, which we treat as an unstable transition state between the PG4 and NPG4 states. Concretely, we define the Gibbs free energies, enthalpies, and entropies, working within the approximation that the formed G4s to have roughly equal enthalpy *H*_0_ and approximating the transition state to have enthalpy *H*^‡^:

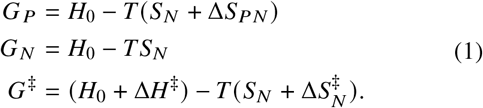

Here, *G* values are Gibbs free energies, *T* is the temperature, *S*_*N*_ is the NPG4 entropy, *S*_*P*_ is the PG4 entropy, 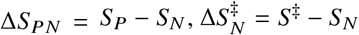 is the entropy difference between the transition state and the NPG4 state, and Δ*H*^‡^ = *H*^‡^ *H*_0_. We take the parallel configuration entropy to be greater than the antiparallel entropy, which makes PG4 thermodynamically more stable than NPG4, consistent with ref. (25). The system can undergo a chemical reaction:

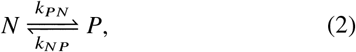

where the scaled reaction rates are Arrhenius-type:

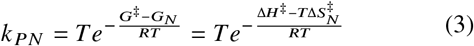

and

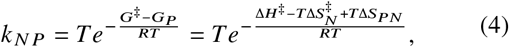

where we have assumed a temperature-proportional Arrhenius constant, as in transition state theory (35), and have divided by the temperature-independent prefactors, writing *R* as the ideal gas constant.

We let *p* (*t*; *T*) be the fraction of PG4s at time *t* and fixed temperature *T*, so 1 − *p* (*t*; *T*) is the fraction of NPG4s. We can write the rate equation

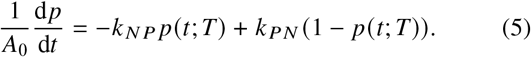

For mathematical convenience, we define a rescaled time *τ* = *A*_0_*t* which now defines the timescale based on the empirical transition attempt frequency. The solution to the above differential equation is

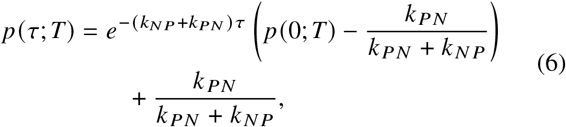

which can be expressed in terms of 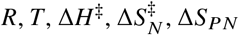, and *τ*.

For some initial fraction *p* (0; *T*) of PG4s, eq. (6) gives the fraction *p* (*τ*; *T*) of PG4s after some fixed scaled time *τ*. In our experiments, temperature is increased periodically, with some fixed wait time between measurements. We assume the the initial PG4 fraction at some new temperature *T*_*i*+1_ in a sequence of temperatures indexed by *i* to be given by the final PG4 fraction at the previous temperature *T*_*i*_. As a result, we can write *p* (0; *T*_*i*_ +_1_) = *p* (*τ*; *T*_*i*_). Now, writing *p*_*i*_ ≡ *p* (*τ*; *T*_*i*_),*K*_*i*_ ≡ *k* _*N P*_ (*T*_*i*_) + *k* _*PN*_ (*T*_*i*_), *r*_*i*_ ≡ *k* _*PN*_ (*T*_*i*_) /*K*_*i*_, and 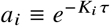, the PG4 fraction at the end of step *i* is concisely

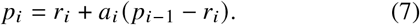

For an algebraic sequence of measurement temperatures *T*_*i*_ = *T*_0_ + *i*Δ*T* (as in our experiments), the parallel fraction after *n* steps can now be written in terms of the initial conditions

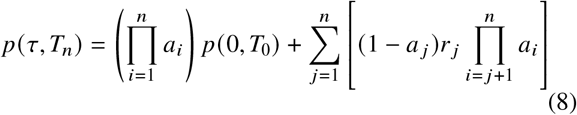

We can then numerically construct a plot of *p* (*τ, T*_*i*_) as a function of temperatures *T*_*i*_. In Figure 5D, we observe that there is a homogenization temperature *T* ^*^ at which the PG4 and NPG4 fractions experience an inflection point. The theory predicts that around this inflection point, the G4 population converts from a heterogeneous population of NPG4s and PG4s to a (nearly) homogeneous population of PG4s. Until this temperature is reached, the fraction of each G4 topology is stable due to kinetic trapping since NPG4s are taken to be a metastable state. The reaction rapidly proceeds after reaching a certain temperature. This is evidence of a *thermally activated* transition of the metastable NPG4s to the stable PG4s. This is not a true thermodynamic phase transition because the ground state configuration does not change; but rather, the theory predicts a kinetic phenomenon where a long-lived metastable state transitions to a stable one when the relative height of the transition state barrier is reduced via heating. By switching the definitions of *G* _*N*_ and *G* _*P*_ in eq. (1), we can see that if PG4s were metastable while NPG4s were globally stable, an increase in temperature would lead to homogenization to NPG4s. This does not agree with experiment and thus suggests that PG4s are more stable than NPG4s for the GGGGCC sequence in the context of our model.

We now build simulated spectra using our derived eq. (8). As we discussed in Section 3.1, hybrid and antiparallel spectra share certain spectroscopic signatures. We posit that, as a result, the hybrid conformation may be difficult to distinguish from a mixture of antiparallel, hybrid, and parallel G4s, so for simplicity we take the parallel and antiparallel structures to be the fundamental “basis spectra” for our theoretical analysis.

CD spectra matching the features of parallel and antiparallel G4s are plotted in Figure 5E. The parallel G4 spectrum, obtained from Figure 2C at 100 °C, closely matches published spectra (31, 36). Since pure antiparallel CD spectra are not available, the antiparallel spectrum is approximated by subtracting off the parallel contribution to an otherwise mixed or hybrid-appearing spectrum at 20 °C. This is accomplished by assuming that at 20 °C there is some fraction *p*_est_ = 0.5 of PG4s and some fraction 1 − *p*_est_ = 0.5 of antiparallel G4s; this choice is arbitrary and is acknolwedged as a limitation of the theory, in addition to the assumption that the spectrum is a linear combination of only two species’ basis spectra. The basis parallel and antiparallel spectra (Figure 5E), *ϵ*_*P*_ (*λ*) and *ϵ* _*A*_ (*λ*), are functions of wavelength *λ* and are used to obtain simulated 3D CD spectra by combining the calculations from the thermodynamic-kinetic theory eq. (8).

Treating the 3D CD spectrum (26, 27) as a linear combination of the basis spectra weighted by the PG4 and NPG4 fractions from eq. (6), we have an expression for the theoretical 3D CD spectrum

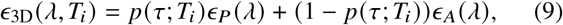

which is plotted as 2D and 3D spectra for various initial PG4 fractions *p* (0; *T*_0_) in Figure 5F-K. The simulated spectra obtained show remarkable qualitative agreement with the experimental 3D and 2D spectra in Figure 2. Changing *p* ( 0; *T*_0_) simply modulates the shape of the initial spectrum at the starting temperature *T*_0_, allowing us to explore heterogeneous initial conditions as we see throughout our experimental datasets (Figure 2, Figure S2, S3, S4, and S5). Changing *p*_est_ affects the 3D CD spectrum by performing an affine shift of the curve *p* (*τ*; *T*_*i*_) (Supplementary Note A).

### Alternative mechanisms

The simulated spectra show that the two-state theory of a metastable-to-stable thermally activated transition is compatible with the experimental results. However, alternative mechanisms may also explain the experimental data. One possibility is that parallel and antiparallel configurations, due to differing thermostability, reversibly melt at different temperatures. As a result, the thermally activated homogenization could actually be melting of antiparallel G4s at lower temperatures subsequent melting of parallel G4s at higher temperatures. This would explain the observed single-sweep experimental data, but would not explain the irreversibility experiments. It is possible, however, that antiparallel G4s could unfold during the first temperature sweep but then find the more stable parallel G4 fold while cooling. This would be similar to ref. (25)’s model that two folding paths exist while folding parallel G4s, and could be implemented mathematically using the model from this paper, with an additional “unfolded” state. Alternatively, the unfolded state or off-target traps could be kinetically trapping unfolded antiparallel G4s and preventing them from refolding. this proposed mechanism would still involve metastability, but in the opposite direction from the one proposed here: metastable trapping happens after heating, not before.

Nucleic acid condensate formation is known to be linked to disease (12, 28–30), it has been shown that GGGGCC RNA repeats indeed undergo liquid-liquid phase separation (12). It is possible that certain topologies are more prone to droplet formation, and phase separation could stabilize particular conformations within a droplet which are not globally stable in the bulk. After a heating cycle, droplet formation itself could become metastable and therefore prevent initial configurational composition from being restored. Thus, aggregation-based mechanisms cannot be ruled out at this time. However, degradation of the DNA can be ruled out as a possible contributing mechanism because we have shown previously that degradation of the nucleic acids does not occur (21).

## 4 DISCUSSION

In this work, we have explored the thermodynamic and kinetic relationships between parallel and non-parallel G4 configurations of the GGGGCC hexanucleotide repeat expansion which appears in ALS/FTD. By performing temperature-swept CD measurements, we plotted 3D CD spectra which showed a clear change in CD spectral shape indicative of homogenization of G4s to the parallel configuration followed by melting of the parallel G4s at subsequently higher temperatures. We then demonstrated experimentally that this transition was irreversible by performing sequential temperature sweeps with a cooling step in between. Finally, we proposed a simple, minimal two-state theory to explain the thermally activated configurational homogenization as a metastable-to-stable transition of kinetically trapped NPG4s into PG4s. We also proposed alternative mechanisms which could involve metastability and kinetic trapping during the refolding process as well as droplet formation and aggregation.

Regardless of the exact theoretical mechanism, our results indicate that temperature-dependent regulation of the free energy landscape governing G4 conformation can lead to substantial changes in the composition of the G4 population. However, we acknowledge a limitation that the broad temperature ranges used *in vitro* in this study are not encountered *in vivo* and therefore may not be directly relevant to cell states. However, since G4s are present *in vivo* (37), it is still possible that intracellular enzymatic regulators could modulate the free energy barriers separating G4 conformations in the way that we used heat to regulate them in our experiments. There are many known helicases which interact with and can unwind G4s such as the DEAH-box helicase, Bloom helicase, and Fanconi anemia complementation group J helicase (38).

More broadly, our work opens the door to further investigation of how G4 configurational composition and thermodynamic/kinetic phenomena can impact DNA synthesis and repair fidelity. Many open questions also remain about how G4 configurational changes may be coupled to biomolecular condensate formation observed in pathological cell states. An understanding of how NPG4-PG4 transitions, unimolecular G4 versus multimolecular G4 formation, and liquid-liquid phase separation are related to each other is still an open question which will be addressed in our forthcoming work.

## 5 AUTHOR CONTRIBUTIONS

D.R., D.E., V.M., and B.K.M. developed the project idea. D.R., O.L., O.M., S.B., S.D., R.B., R.D., and B.K.M. conducted experiments. V.M. contributed analytical theory. D.R., V.M., and B.K.M. analyzed the data. D.R., V.M., and B.K.M. wrote the manuscript. B.K.M. supervised the project.

## 6 ACKNOWLEDGMENTS

## Funding and resource acknowledgment

This work was supported by VCOM’s REAP grants 1032453 and 1302559 (to B.K.M.), a Hertz Foundation Fellowship (to V.M.), a PD Soros Fellowship (to V.M.), and by award T32GM14427 from the National Institute of General Medical Sciences (to Harvard/MIT MD-PhD Program). The content is solely the responsibility of the authors and does not necessarily represent the official views of the National Institute of General Medical Sciences or the National Institutes of Health. The authors declare no known conflict of interest. The authors thank three anonymous reviewers for their helpful comments and suggestions.

## Code and data availability

Code and data will be made publicly available upon publication.

## SUPPLEMENTARY INFORMATION

### Supplementary Note A MODULATING *p*_est_ PERFORMS AN AFFINE SHIFT ON *p* (*τ, T*_*i*_)

Suppose the experimental CD specturm at 20 °C, *ϵ*_mix_(*λ*) is given by a linear combination of parallel and antiparallel basis spectra, *ϵ*_*P*_ (*λ*) and *ϵ* _*A*_(*λ*), respectively, with a guessed PG4 fraction *p*_est_ and NPG4 fraction 1 − *p*_est_:

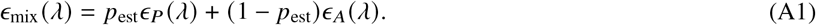

The estimated antiparallel basis spectrum is solved in terms of the experimentally observed room temperature (mixed) spectrum *ϵ*_mix_(*λ*) and the presumed parallel spectrum (obtained at high temperature) *ϵ*_*P*_ (*λ*):

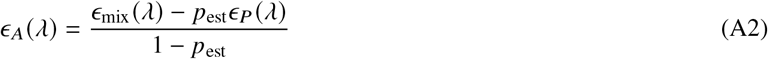

The theoretical 3D spectrum *ϵ*_3D_(*λ, T*_*i*_) is given by main text eq. (6). Substituting the above equation for *ϵ* _*A*_(*λ*), we have

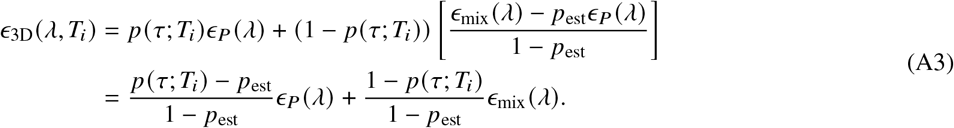

Letting

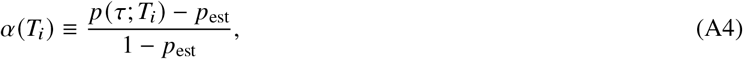

we can now write

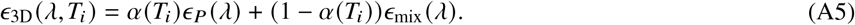

Thus, the 3D CD spectrum is only affected by changes to *α* (*T*_*i*_), which is a linear function of the *p* (*τ*; *T*_*i*_) given by the thermodynamic-kinetic theory in the main text. If we choose a different *p*_est_, it simply affects via an affine shift of the curve *p* (*τ*; *T*_*i*_).

### SUPPLEMENTARY TABLE

**Table S1:** Summary of all 3D CD spectra obtained, their *T*_peak_ values, and their locations within this paper.

| Number of Repeats | Replicate Number | $T_{\text{peak}}$ (°C) | Figure | Additional conditions |
| --- | --- | --- | --- | --- |
| 2 | 1 | 81.98 | S2 |  |
| 4 | 1 | 86 | 2, repeated in S2 |  |
| 4 | 2 | 94.02 | S4 |  |
| 4 | 3 | 91.98 | S4 | 300 second wait time |
| 6 | 1 | 81.99 | S2 |  |
| 6 | 2 | 79.99 | S3 |  |
| 10 | 1 | 88.02 | S2 |  |
| 14 | 1 | 84.02 | 2, repeated in S2 |  |
| 14 | 2 | 84 | S3 |  |
| 16 | 1 | 84.02 | S2 |  |
| 16 | 2 | 83.99 | S5 | 300 second wait time |
| 16 | 3 | 84 | 2 | with PEG, extended temperature range |
| 17 | 1 | 82.02 | S2 |  |
| 18 | 1 | 84.01 | S2 |  |
| 20 | 1 | 83.98 | 2, repeated in S2 |  |

### SUPPLEMENTARY FIGURES

**Figure S1:**
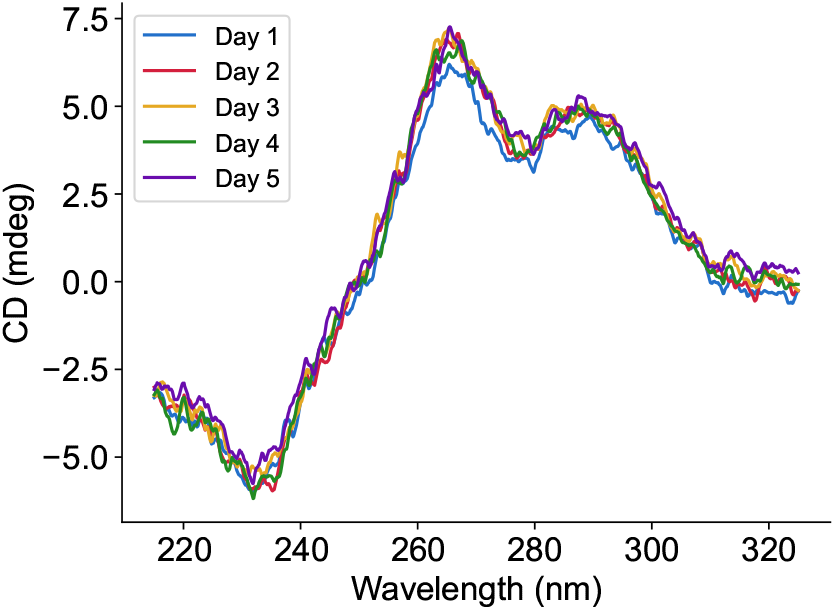
For 17 repeats of GGGGCC, the CD spectrum remained the same and stable at room temperature each day, for 5 days.

**Figure S2:**
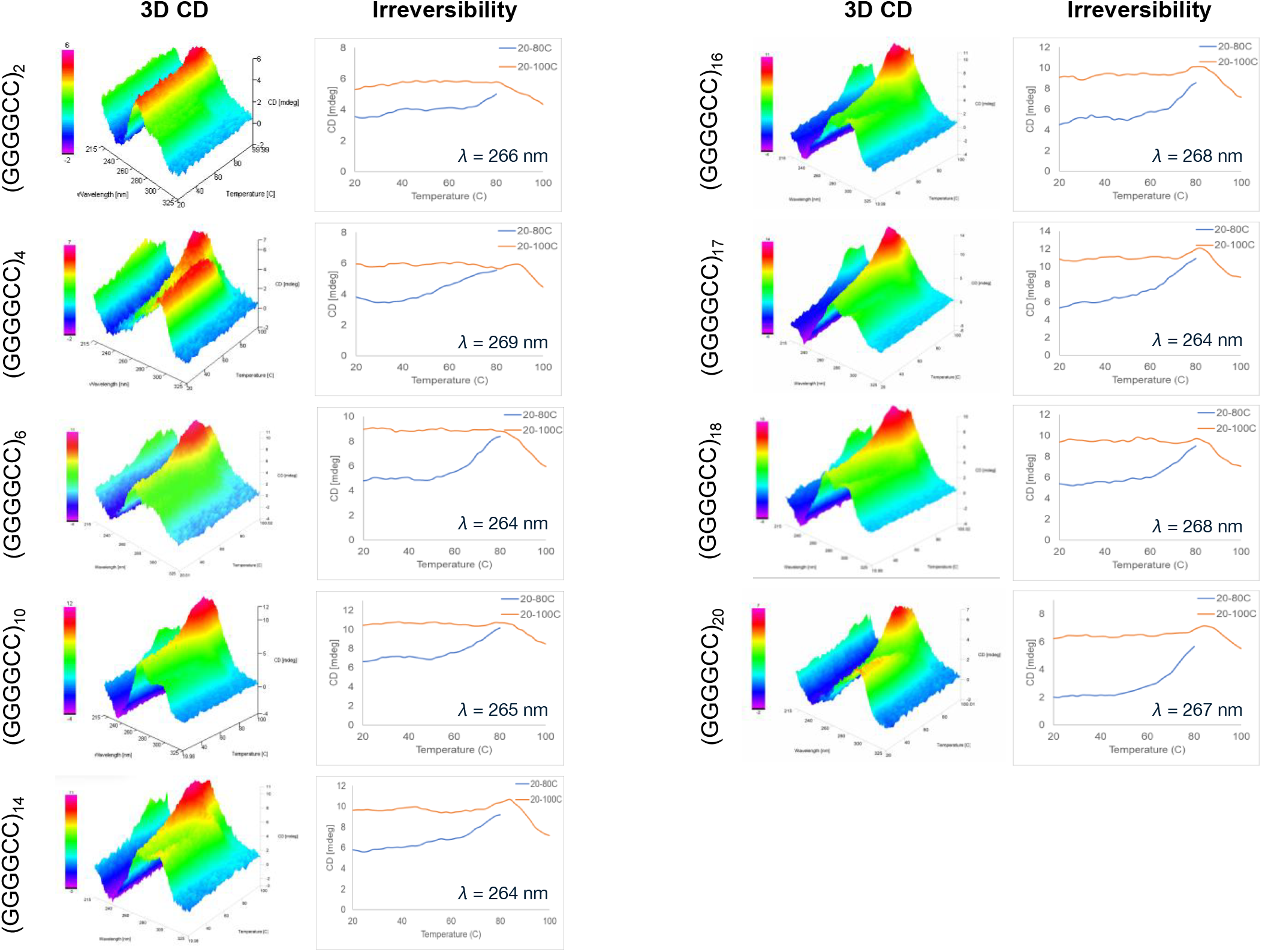
3D CD spectra and irreversibility data for additional oligonucleotides with varying hexanucleotide repeat copy numbers. 3D CD spectra demonstrate the metastable-to-stable homogenization transition consistent with theory, and consistent across hexanucleotide repeat copy numbers. Irreversibility data show stability of the CD peak during the second temperature sweep (20 °C to 100 °C, orange) after it was reached at the end of the first temperature sweep (20 °C to 80 °C, blue). Irreversibility data are consistent across hexanucleotide repeat copy numbers. The wavelengths used to capture irreversibility data for each copy number are listed and are all in the 264-269 nm.

**Figure S3:**
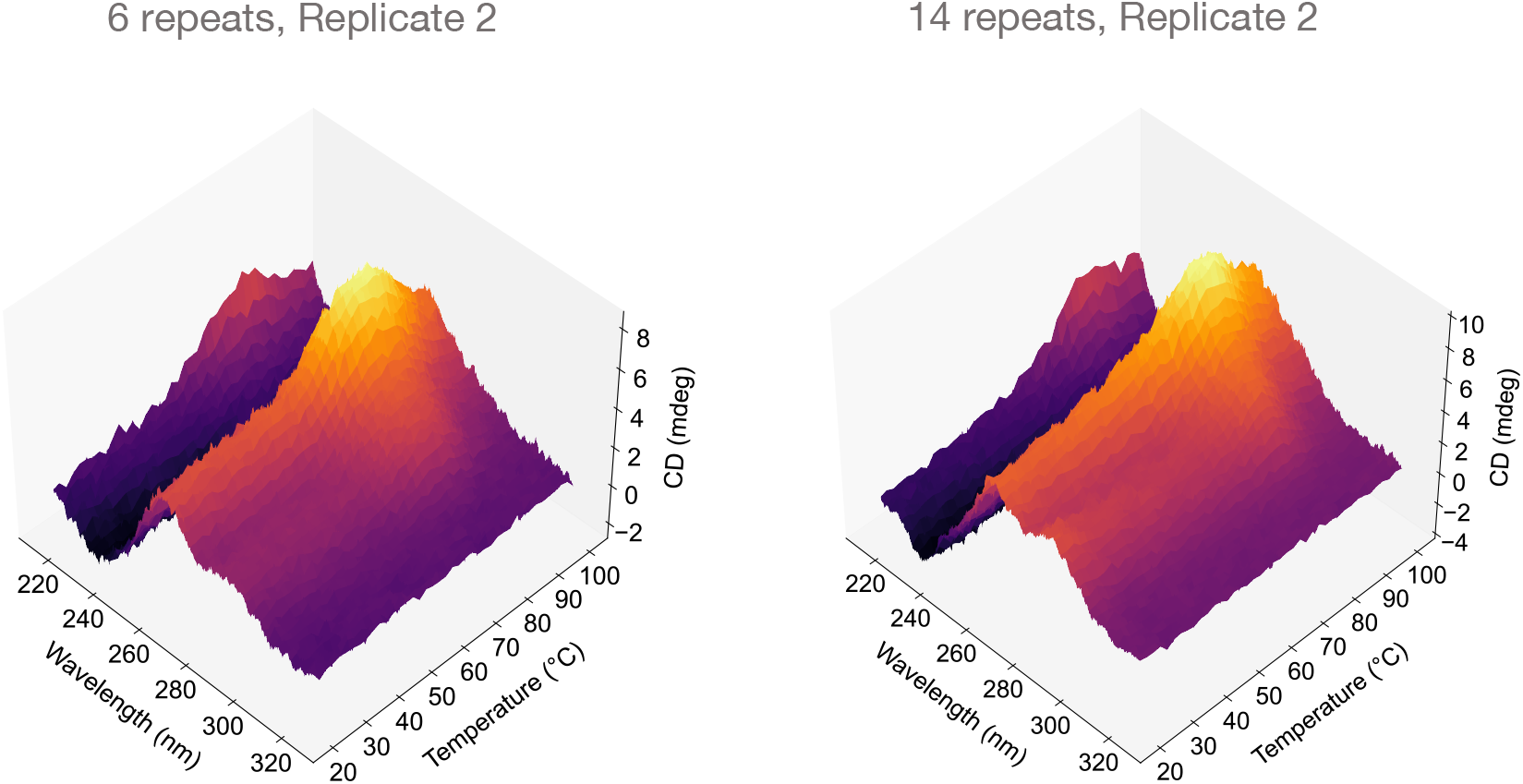
Additional experimental replicates for 6 and 14 repeats not utilized in other figures. Results are qualitatively similar to Figure S2.

**Figure S4:**
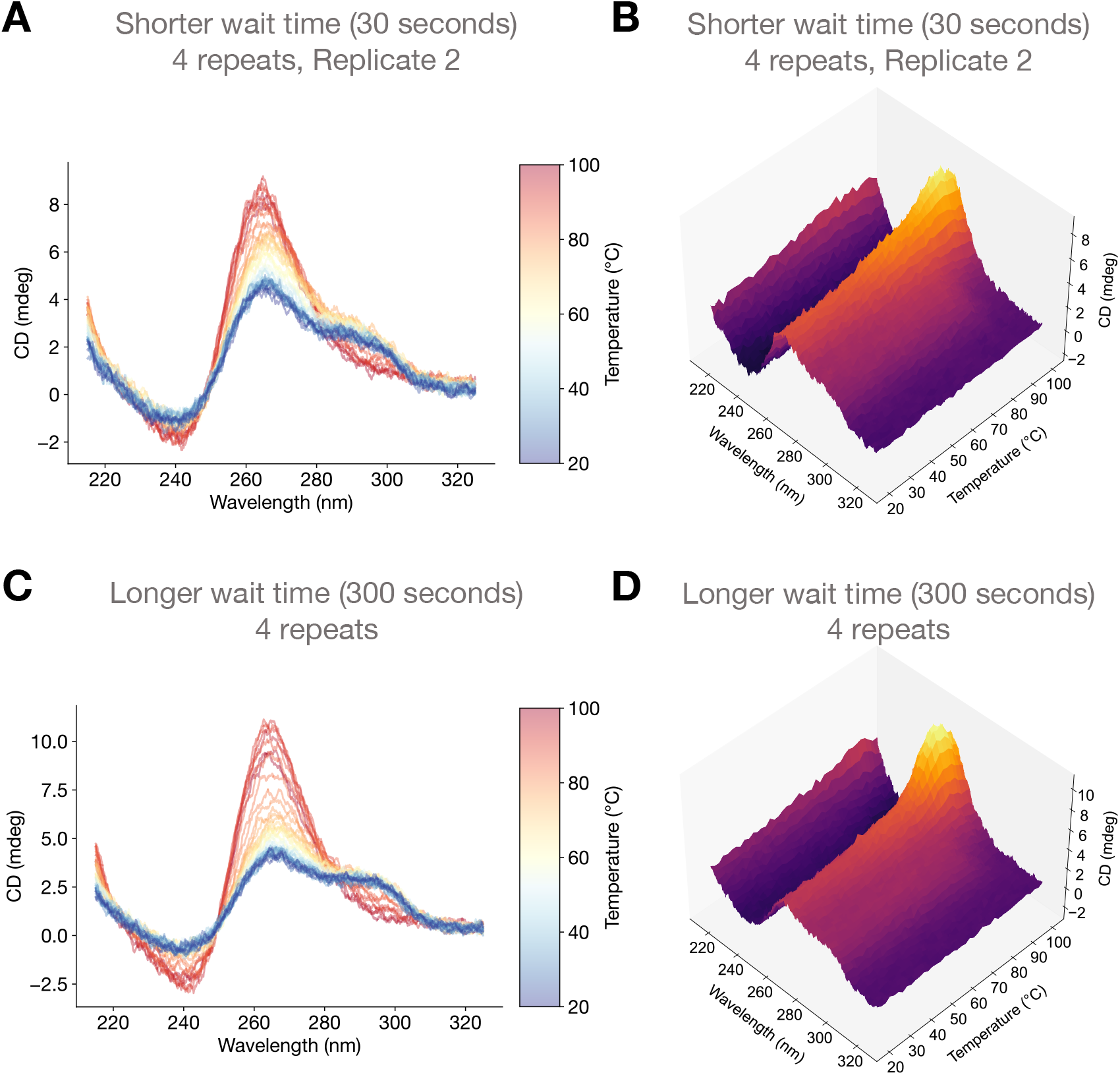
For 4 repeats of GGGGCC, we plot the temperature sweep experiments with (**A**,**C**) color-coded 2D and (**B**,**D**) 3D spectra with both shorter (30 second) and longer (300 second) wait times between each 2 °C increment.

**Figure S5:**
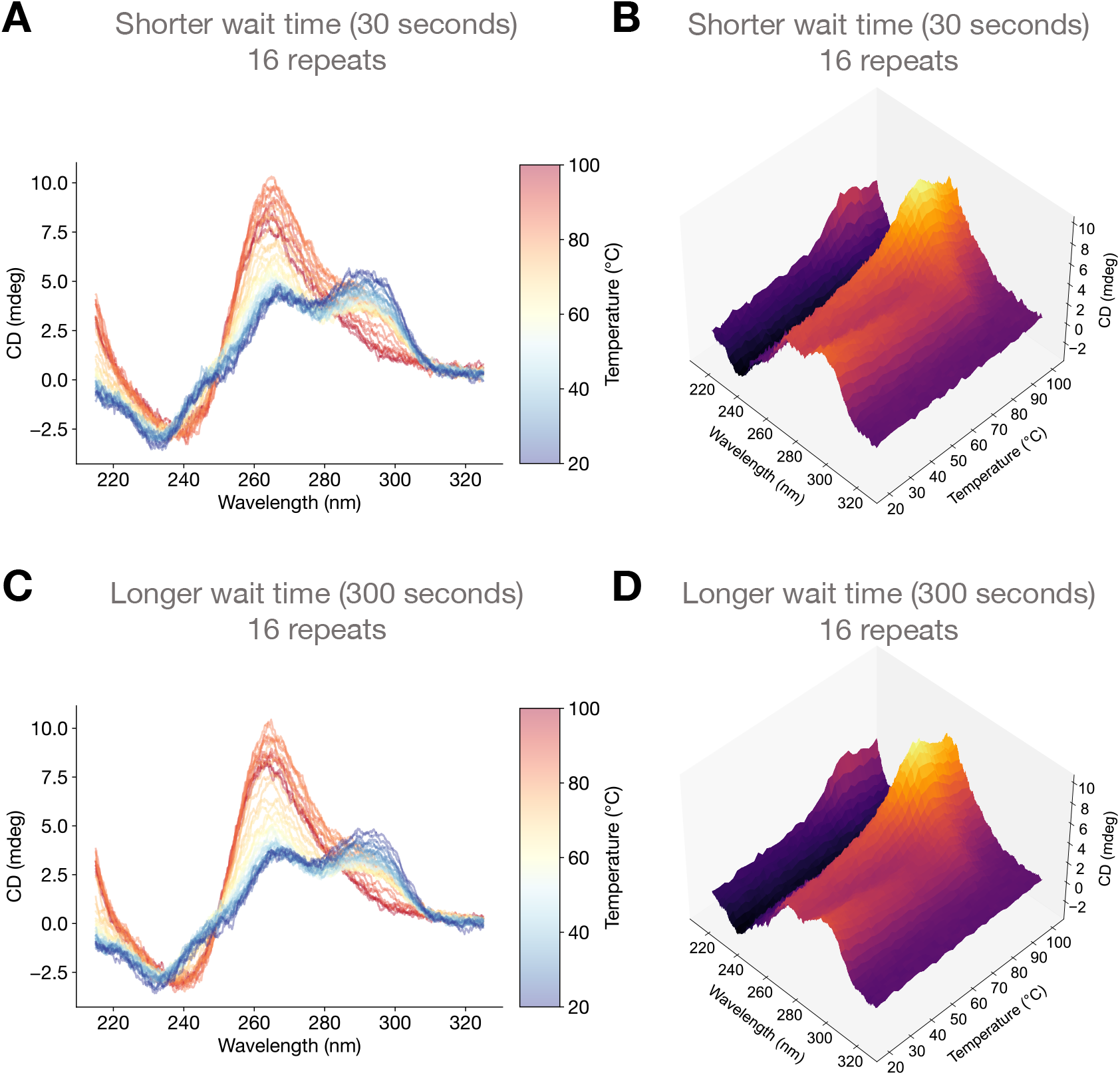
For 16 repeats of GGGGCC, the temperature sweep experiments yielded similar (**A**,**C**) color-coded 2D and (**B**,**D**) 3D spectra with both shorter (30 second) and longer (300 second) wait times between each 2 °C increment.

**Figure S6:**
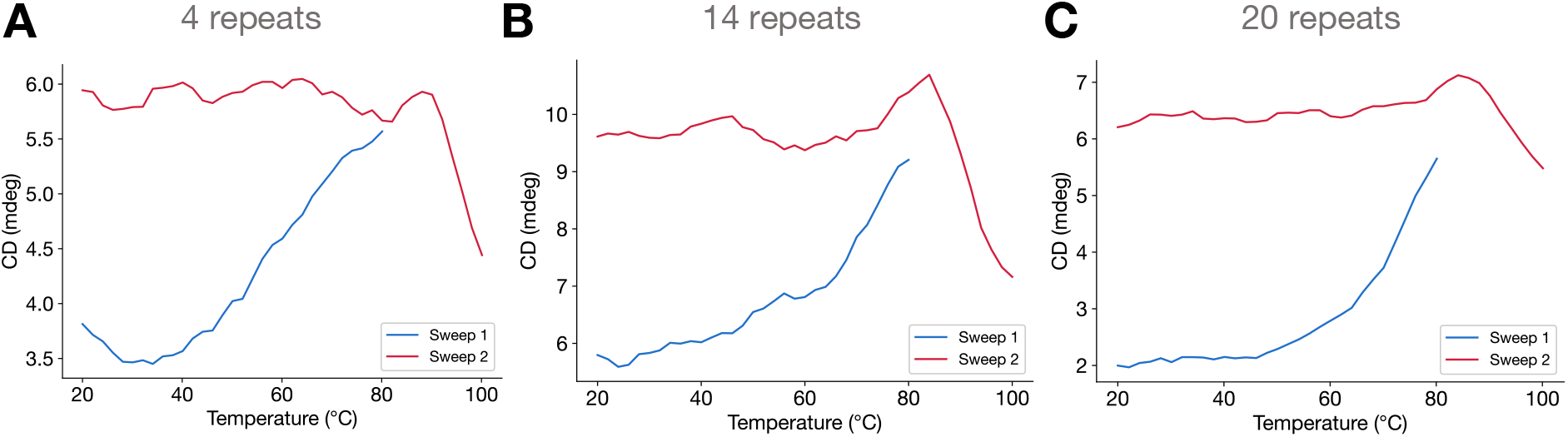
Thermally activated metastable-to-stable G4 transition is irreversible. (**A**) For 4 hexanucleotide repeats, the first temperature sweep (blue) shows a CD increase at 269 nm as temperature increases from 20 °C up to 80 °C, indicative of a transition to PG4s. After cooling back to 20 °C, the second temperature sweep (red) shows stability of the CD peak from 20 °C up to roughly 80 °C, after which CD decreases, either due to G4 melting (or possibly NPG4 formation). (**B**) Same as (A), but with 14 hexanucleotide repeats with CD measured at 264 nm. (**C**) Same as (A) and (B), but with 20 hexanucleotide repeats with CD measured at 267 nm. CD lines were smoothed with a moving average filter using a convolution kernel with a 5-datapoint width.

